# Thoracoabdominal pressure transmission during prone and supine cardiopulmonary resuscitation in fresh-frozen human cadavers

**DOI:** 10.64898/2026.09.22.753658

**Authors:** Zachary Smith, Robert Lust, Christian Falyar, Mel Swanson, Bimbola F. Akintade, Linda Bolin

**Affiliations:** College of Nursing, East Carolina University, 2205 W 5th St, Greenville, North Carolina 27834, USA; Department of Physiology, Brody School of Medicine, East Carolina University, 600 Moye Blvd, Greenville, North Carolina 27834, USA; Middle Tennessee School of Anesthesia, 315 Hospital Drive, Madison, Tennessee 37115, USA

**Keywords:** resuscitation physiology, prone positioning, pressure transmission, thoracic pump, abdominal counterpressure, cadaveric model, chest compression

## Abstract

**Aim:** Prone cardiopulmonary resuscitation (CPR) may be necessary when turning a prone patient supine would delay chest compressions. Although prone compressions can generate arterial pressures comparable with or greater than supine CPR, the pathway of pressure transmission is uncertain. We examined synchronized intrathoracic, intra-abdominal, and central arterial pressures in both supine and prone positions.

**Methods:** Two thawed fresh-frozen adult cadavers underwent three, 2-minute mechanical CPR trials per position in a counterbalanced crossover sequence. Solid-state catheters recorded pleural, peritoneal, and central arterial pressures simultaneously. Trial-level outcomes included peak pressure, mean pressure, pressure-time area, and the mean peritoneal-to-pleural pressure gradient. Exploratory fixed-effects models included position, cadaver, and their interaction.

**Results:** Prone CPR increased peak intrathoracic pressure by 7.04 mmHg, peak intra-abdominal pressure by 21.69 mmHg, and peak arterial pressure by 15.40 mmHg. Mean intra-abdominal and arterial pressures increased by 16.22 and 9.90 mmHg, respectively. The mean peritoneal-to-pleural gradient reversed direction from −8.46 mmHg supine to 4.85 mmHg prone. Intrathoracic pressure-time area increased 3.4-fold, from 1.62 to 5.46 mmHg·s, and arterial pressure-time area increased 2.2-fold, from 2.96 to 6.42 mmHg·s.

**Conclusion:** Compared to supine, prone mechanical CPR generated higher arterial pressures and altered the pressure relationship across the thoracoabdominal boundary in both cadavers. Higher abdominal pressure coincided with a larger intrathoracic pressure-time area, a pattern compatible with reduced caudal pressure dissipation. Because diaphragm motion was not measured, this proposed mechanism remains inferential.

## Introduction

Prone cardiac arrest is reported most often in operating rooms and intensive care settings, where airway devices, vascular access, surgical equipment, and an open operative field can complicate repositioning [1–4]. Current American Heart Association guidance states that prone CPR may be considered when a patient cannot be safely turned supine [5]. The European Resuscitation Council similarly recommends initiating CPR in the prone position when cardiac arrest occurs with an advanced airway already in place [6]. In simulation, initiating compressions in the prone position reduced time to first compression and increased chest compression fraction compared with turning the patient supine before CPR [7].

Clinical and experimental reports suggest that prone compressions can generate arterial pressures comparable with or greater than those produced during supine CPR [1–4,8,9]. Most studies have focused on feasibility, isolated pressure observations, or clinical outcomes. To our knowledge, none has simultaneously measured pressure changes in the abdomen, thorax, and central arterial system. The pathway from posterior chest compression to generated arterial pressure therefore remains uncertain.

Closed-chest CPR is commonly explained by cardiac pump and thoracic pump mechanisms [10–13]. Abdominal pressure may also influence venous return, arterial pressure, and recoil-phase pressure relationships [13,14]. Prone positioning changes contact among the abdomen, thorax, support surface, and compression device [15,16]. Abdominal counterpressure may limit caudal pressure dissipation across the diaphragm and allow a larger or more sustained pressure pulse within the thorax. This mechanism has not been tested directly in a human cadaver model.

This pilot study compared synchronized intrathoracic, intra-abdominal, and central arterial pressures during prone and supine mechanical CPR in two fresh-frozen human cadavers. We examined whether higher arterial pressure during prone CPR was accompanied by higher intra-abdominal pressure, a shift in the peritoneal-to-pleural pressure gradient, and a larger intrathoracic pressure waveform.

## Methods

### Study design and setting

A prospective within-specimen crossover study was performed in the cadaver laboratory at the Middle Tennessee School of Anesthesia in Madison, Tennessee. Two thawed fresh-frozen adult human cadavers were obtained through the Medical Education and Research Institute. Each cadaver underwent mechanical CPR in both positions and served as its own anatomic control. Position order was counterbalanced between cadavers to reduce potential sequential order effects. Cadaver A underwent supine CPR followed by prone CPR. Cadaver B underwent prone CPR followed by supine CPR.

Cadaver A was a male donor aged ≥90 years with a height of 168 cm, weight of 54 kg, body mass index of 19.2 kg/m², and a history of coronary stents. Cadaver B was a female donor aged 87 years, with a height of 160 cm, weight of 57 kg, body mass index of 22.3 kg/m², with a history of coronary stents, and a prior hip replacement. Both cadavers had intact thoracic and abdominal anatomy suitable for mechanical compression and invasive pressure monitoring.

### Vascular preparation and instrumentation

Before laboratory instrumentation, a licensed mortician performed vascular conditioning to reduce arterial clot burden and improve pressure transmission. The preparation was adapted from a prior human cadaver CPR model [17]. Metaflow Vascular Conditioner (The Dodge Company, Billerica, Massachusetts, USA) was mixed with 0.9% saline and delivered through the right common carotid artery with an APC24 mortuary perfusion machine (The Dodge Company) in pulsatile mode. Controlled drainage was established through the right internal jugular vein. The right neck vessels were ligated after conditioning.

The trachea was intubated with an 8.0 endotracheal tube, which was clamped during all compression trials to standardize the thoracic pressure environment. Central arterial pressure was measured with a Millar SPR-524 Mikro-Tip catheter (Millar, Pearland, Texas, USA) introduced through a left common carotid cutdown approach and advanced toward the aortic arch. Intrathoracic pressure was measured from the left pleural space through a Veress-assisted introducer at the left second intercostal space. Intra-abdominal pressure was measured through a left periumbilical peritoneal introducer. Guidewire position was confirmed with ultrasound at the carotid, pleural, and peritoneal access sites before sheath or catheter advancement.

### Pressure acquisition and calibration

The three pressure signals at the access sites were acquired simultaneously with a PowerLab system, an FE224 Quad Bridge Amplifier, and LabChart software (ADInstruments, Colorado Springs, Colorado, USA). Each catheter remained assigned to the same monitoring channel throughout the study. Prior to insertion, each pressure catheter underwent a two-point unit conversion with an MLA6595 Delta-Cal pressure transducer simulator. Calibration points were set at 40 and 80 mmHg. After catheter placement, the investigators confirmed a stable baseline along with a visible waveform response with test compressions. Signals showed no clipping or excessive electrical noise before formal trials began.

### Mechanical CPR protocol

Mechanical compressions were performed with a LUCAS 3 Chest Compression System from Stryker (Portage, Michigan, USA). For the purposes of this study, the term CPR refers specifically to mechanical chest compressions, as ventilation was not administered during the trials. The LUCAS 3 was operated in continuous mode at a programmed rate of approximately 102 compressions/min and a nominal compression depth of 5.3 cm. Delivered rate and depth were not independently measured during testing. For supine trials, the suction cup was centered over the lower half of the sternum in accordance with adult CPR guidance [5]. For prone trials, it was centered over the T7-T9 region near the inferior scapular level and posterior projection of the heart, in line with locations described in prior prone CPR reports [1–4,8,9].

Each position began with a 5-minute stabilization period. Three formal trials were then performed. Each trial included a 20-second compression stabilization interval followed by 2 minutes of recorded mechanical CPR. A 1-minute pause separated trials. After repositioning, the cadaver was rested for 5 minutes before the sequence was repeated in the opposite position.

### Outcome definitions and waveform processing

Trial-level averages were used as the primary unit of analysis. For each 2-minute trial, peak and mean pressures were summarized for the intrathoracic, intra-abdominal, and arterial channels. LabChart Peak Analysis calculated pressure-time area for each included compression cycle, and cycle-level values were averaged within the trial. The mean peritoneal-to-pleural pressure gradient was calculated at the trial level as mean intra-abdominal pressure minus mean intrathoracic pressure. Negative values indicated higher mean intrathoracic pressure, where positive values indicated higher mean intra-abdominal pressure.

Composite waveforms were generated in LabChart separately for each cadaver and body position. Using LabChart Scopeview, waveforms were aligned and averaged using all included compression cycles from all three trials in each position. Each composite represents the mean pressure waveform for one compression cycle rather than a manually selected or superimposed beat.

### Statistical analysis

Continuous outcomes are reported as mean and standard deviation. Each outcome was analyzed using a two-factor, fixed-effects general linear model that included body position, cadaver, and the position-by-cadaver interaction. Estimated marginal means were used to calculate the prone minus supine difference and 95% confidence interval. F statistics, p values, and partial eta squared are reported for the position effect and the interaction. Analyses were performed in IBM SPSS Statistics for Windows, version 30.0 (IBM Corp., Armonk, New York, USA).

The dataset contained 12 trial-level averages. The three trials within each cadaver and position were repeated technical observations, not independent biological specimens. Because only two cadavers were studied, inferential analyses were considered exploratory. Interpretation emphasized effect estimates, confidence intervals, the direction of change within each cadaver, and position-by-cadaver differences.

### Ethics

No living human participants were enrolled. The University and Medical Center Institutional Review Board (UMCIRB) determined that the project was not human subjects research as it involved deceased human cadaveric donors. Consent for anatomical donation and permitted research or education use was managed by the body donation service before specimen release. Donor identifiers were not included in the study dataset or manuscript.

## Results

### Trial completion

All 12 formal trials were completed with interpretable intrathoracic, intra-abdominal, and arterial waveforms. The analysis included three supine and three prone trial averages from each cadaver. Table 1 presents cadaver-specific descriptive statistics. Table 2 presents the overall position effects and secondary outcomes.

**Table 1.** Pressure measures by cadaver and cardiopulmonary resuscitation position.

| **Measure** | **Cadaver A** |  | **Cadaver B** |  |
| --- | --- | --- | --- | --- |
|  | **Supine mean (SD)** | **Prone mean (SD)** | **Supine mean (SD)** | **Prone mean (SD)** |
| ITP peak pressure | 26.53 (0.25) | 33.77 (2.04) | 32.69 (0.71) | 39.53 (0.44) |
| ITP mean pressure | 9.86 (0.70) | 15.87 (1.66) | 15.95 (0.18) | 15.77 (1.43) |
| IAP peak pressure | 15.39 (0.42) | 31.19 (0.66) | 9.62 (0.29) | 37.21 (1.83) |
| IAP mean pressure | 5.83 (0.50) | 17.88 (1.76) | 3.07 (0.25) | 23.47 (2.94) |
| Arterial peak pressure | 25.97 (1.80) | 42.61 (3.71) | 33.86 (1.03) | 48.02 (2.07) |
| Arterial mean pressure | 7.98 (1.03) | 21.21 (3.22) | 14.03 (0.20) | 20.60 (1.32) |

*Note.* All pressure values are in mmHg. Each cell is based on three trial-level averages. ITP is intrathoracic pressure. IAP is intra-abdominal pressure. SD is standard deviation.

**Table 2.** Overall differences between prone and supine cardiopulmonary resuscitation.

| **Outcome** | **Supine mean (SD)** | **Prone mean (SD)** | **Mean Difference**<br>**(95% CI)** | **F(1, 8)** | **p** | **Partial**<br>**eta**<br>**squared** |
| --- | --- | --- | --- | --- | --- | --- |
| Peak ITP | 29.61 (3.40) | 36.65 (3.42) | 7.04 (5.57-8.52) | 121.44 | < .001 | .94 |
| Peak IAP | 12.51 (3.18) | 34.20 (3.52) | 21.69 (20.35-23.03) | 1392.49 | < .001 | .99 |
| Peak arterial pressure | 29.92 (4.52) | 45.31 (4.00) | 15.40 (12.25-18.55) | 127.16 | < .001 | .94 |
| Mean ITP | 12.91 (3.36) | 15.82 (1.39) | 2.91 (1.38-4.45) | 19.18 | .002 | .71 |
| Mean IAP | 4.45 (1.55) | 20.67 (3.75) | 16.22 (13.91-18.54) | 261.89 | < .001 | .97 |
| Mean arterial pressure | 11.00 (3.38) | 20.90 (2.23) | 9.90 (7.48-12.32) | 89.00 | < .001 | .92 |
| Mean IAP - ITP gradient | -8.46 (4.85) | 4.85 (3.30) | 13.31 (12.17-14.46) | 720.26 | < .001 | .99 |
| ITP pressure-time area | 1.62 (0.40) | 5.46 (0.81) | 3.85 (2.98-4.72) | 103.92 | < .001 | .93 |
| IAP pressure-time area | 1.87 (0.87) | 3.93 (1.23) | 2.06 (0.72-3.39) | 12.66 | .007 | .61 |
| Arterial pressure-time area | 2.96 (0.50) | 6.42 (0.99) | 3.45 (2.40-4.50) | 57.01 | < .001 | .88 |

*Note.* Means and standard deviations are based on six trial-level averages per position. Differences are prone minus supine. Pressure and gradient values are in mmHg. Pressure-time area values are in mmHg·s. F tests are the overall position effect from the two-factor fixed-effects model. ITP is intrathoracic pressure. IAP is intra-abdominal pressure. CI is confidence interval.

### Peak and mean pressure

Peak pressure was higher during prone CPR in all three compartments and in both cadavers (Fig. 1A). Peak intrathoracic, intra-abdominal, and arterial pressures increased by 7.04, 21.69, and 15.40 mmHg, respectively (Table 2). Peak intrathoracic and arterial increases were similar between cadavers. The intra-abdominal increase was larger in Cadaver B than in Cadaver A, at 27.59 and 15.80 mmHg, respectively.

**Fig. 1.**
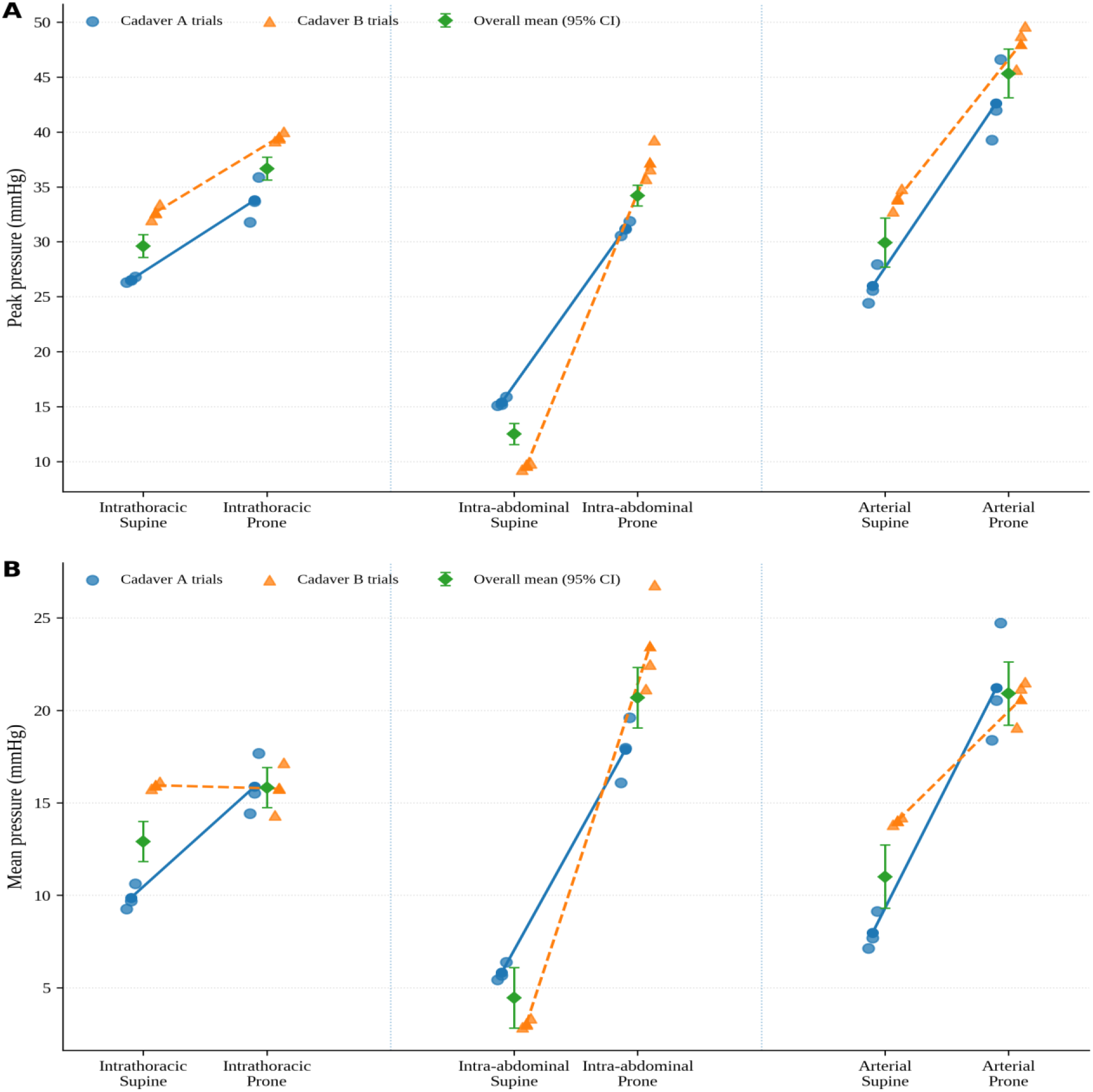
Peak and mean pressures by cardiopulmonary resuscitation position. Panels show (A) peak pressure and (B) mean pressure for the intrathoracic, intra-abdominal, and arterial channels. Small points are trial-level averages, with three trials in each cadaver and position cell. Circles and solid lines are Cadaver A means. Triangles and dashed lines are Cadaver B means. Diamonds and error bars are overall estimated marginal means with model-based 95% confidence intervals.

Mean intra-abdominal and arterial pressures were also higher during prone CPR in both cadavers (Fig. 1B). The overall differences were 16.22 and 9.90 mmHg, respectively (Table 2). Mean intrathoracic pressure differed by cadaver. It increased by 6.01 mmHg in Cadaver A and changed by −0.18 mmHg in Cadaver B.

### Thoracoabdominal pressure gradient

The mean peritoneal-to-pleural pressure gradient reversed between positions in both cadavers (Fig. 2). Mean intrathoracic pressure exceeded mean intra-abdominal pressure during supine CPR, yielding a gradient of −8.46 mmHg. During prone CPR, mean intra-abdominal pressure exceeded mean intrathoracic pressure, yielding a gradient of 4.85 mmHg. The overall shift was 13.31 mmHg and was larger in Cadaver B than in Cadaver A, at 20.58 and 6.04 mmHg, respectively (Table 2).

**Fig. 2.**
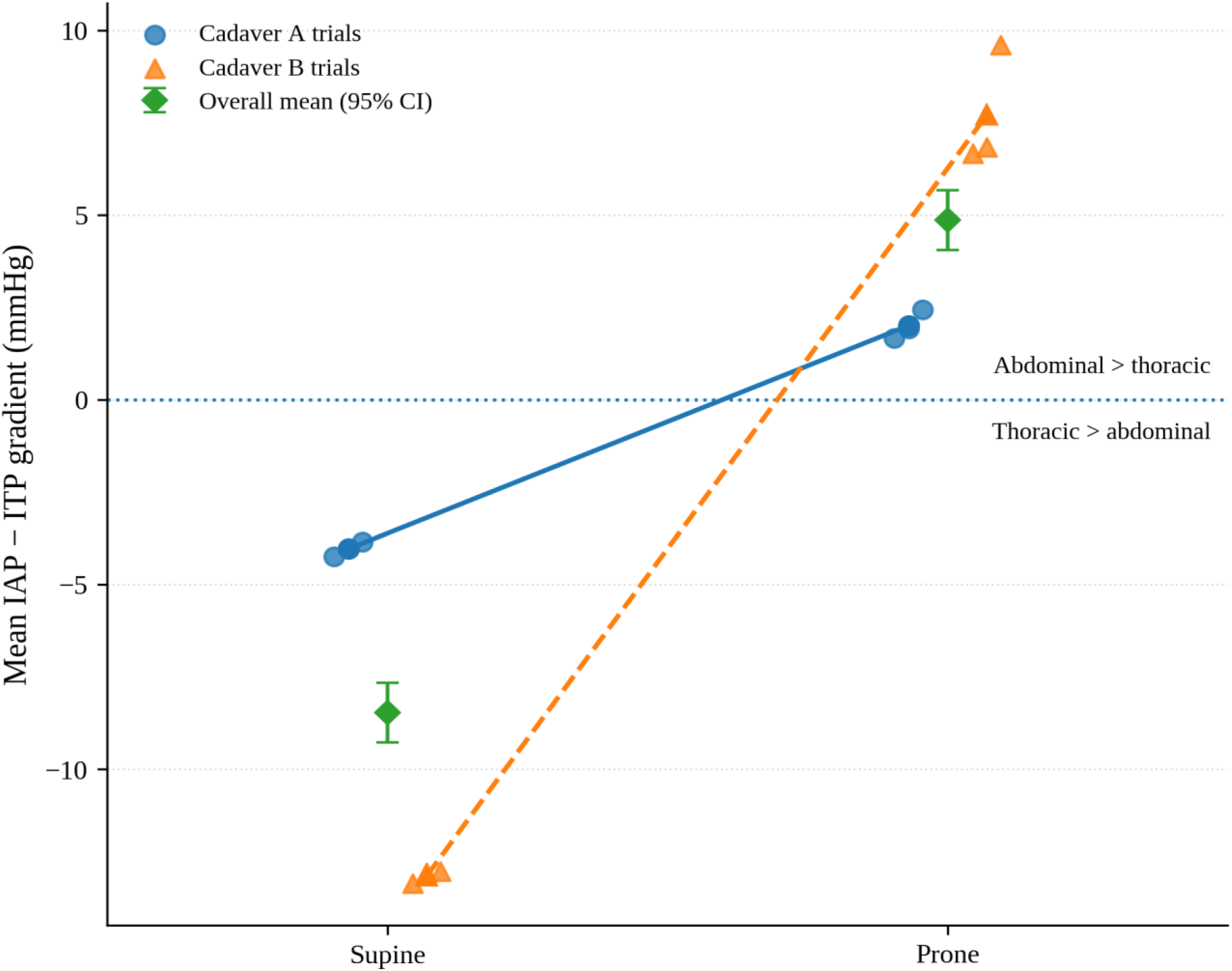
Mean peritoneal-to-pleural pressure gradient by cardiopulmonary resuscitation position. The gradient was calculated as mean intra-abdominal pressure minus mean intrathoracic pressure. The horizontal reference at 0 mmHg indicates equal mean pressure across the thoracoabdominal boundary. Small points are trial-level averages. Circles and solid lines are Cadaver A means. Triangles and dashed lines are Cadaver B means. Diamonds and error bars are overall estimated marginal means with model-based 95% confidence intervals.

### Pressure-time area and composite waveforms

Pressure-time area was greater during prone CPR in all three compartments and in both cadavers (Fig. 3). Intrathoracic, intra-abdominal, and arterial pressure-time areas increased by 3.85, 2.06, and 3.45 mmHg·s, respectively (Table 2). The direction of change was the same in both cadavers for all three pressure-time area measures.

**Fig. 3.**
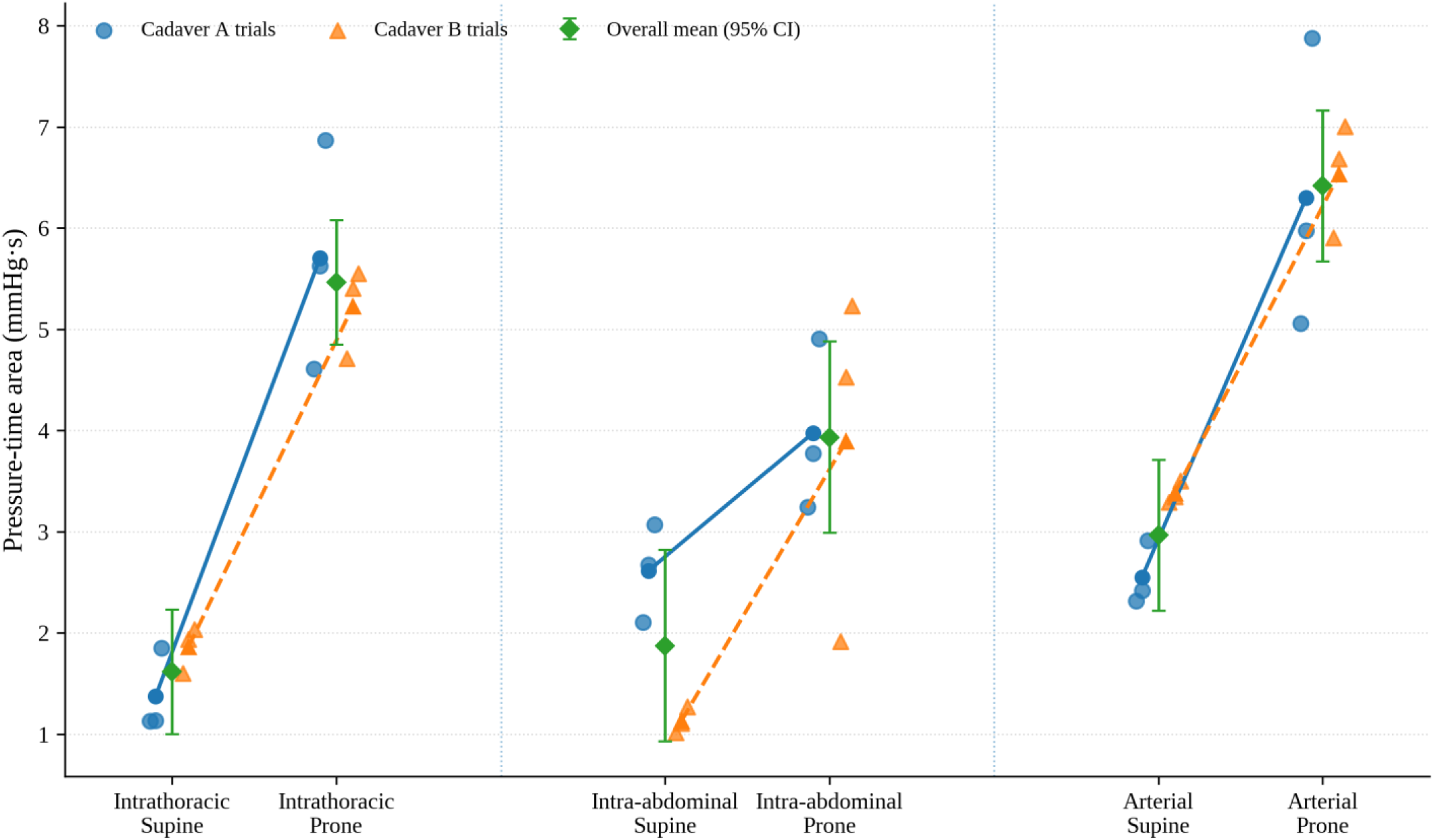
Pressure-time area by cardiopulmonary resuscitation position and pressure compartment. Small points are trial-level averages, with three trials in each cadaver and position cell. Circles and solid lines are Cadaver A means. Triangles and dashed lines are Cadaver B means. Diamonds and error bars are overall estimated marginal means with model-based 95% confidence intervals.

Composite pressure waveforms are shown in Fig. 4. The prone traces had higher peaks and broader width in the intrathoracic, intra-abdominal, and arterial channels. This pattern was consistent with the higher peak pressure and pressure-time area measured during prone CPR.

**Fig. 4.**
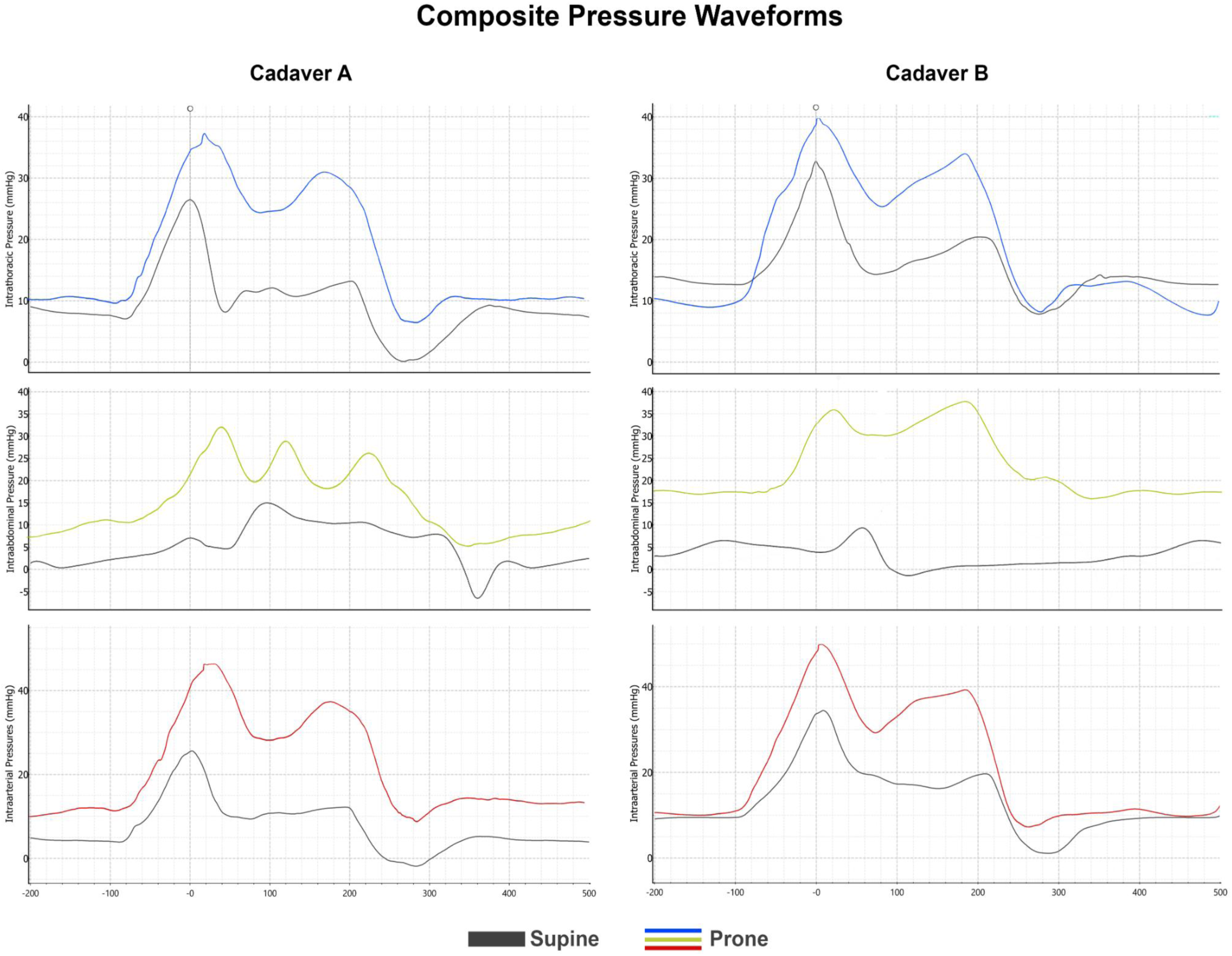
Composite pressure waveforms during prone and supine cardiopulmonary resuscitation. Composite waveforms are shown for Cadaver A and Cadaver B. Each trace is the LabChart-generated average of every included compression cycle across the three trials for one body position. Gray traces are supine. Blue, yellow, and red traces are prone intrathoracic, intra-abdominal, and arterial waveforms, respectively. The x-axis shows time in milliseconds relative to the aligned supine compression cycle. The composite traces illustrate the higher peaks and broader pressure contours measured during prone CPR.

## Discussion

### Primary findings

Prone mechanical CPR generated higher central arterial pressure than supine CPR in both cadavers. The peak arterial increase was similar between specimens, and arterial pressure-time area increased in both. This difference extended across more of the compression cycle rather than being confined to a brief maximum. These arterial changes occurred with higher peak intrathoracic pressure, markedly higher intra-abdominal pressure, and reversal of the mean peritoneal-to-pleural pressure gradient.

### Thoracoabdominal pressure transmission

The gradient between the thoracic and abdominal cavities provides the most direct pressure-based evidence for the proposed thoracoabdominal mechanism. During supine CPR, mean intrathoracic pressure exceeded mean intra-abdominal pressure. During prone CPR, the relationship reversed in both cadavers. The abdomen presented a markedly different pressure environment on the caudal side of the diaphragm in the prone position.

Contact between the anterior abdomen and the support surface may have created counterpressure during posterior thoracic compression. Higher pressure below the diaphragm may limit caudal displacement or reduce dissipation of compression pressure into the abdomen.

The larger intrathoracic pressure-time area in both cadavers suggests that the thoracic pressure pulse was greater and persisted longer. The parallel change in the arterial waveform is consistent with transmission of that pressure pulse into the central arterial system.

The pressure gradient does not establish diaphragm motion, but it changed in the direction expected if caudal pressure dissipation were reduced.

### Peak and mean intrathoracic pressure

Peak intrathoracic pressure increased by a similar amount in both cadavers. Cadaver A had a clear increase in mean intrathoracic pressure. In Cadaver B, mean intrathoracic pressure was nearly unchanged despite increases in peak intrathoracic pressure, intra-abdominal pressure, arterial pressure, and pressure-time area. Cycle mean intrathoracic pressure alone may therefore be insufficient to describe the pressure transmission associated with higher arterial pressure.

Peak pressure and pressure-time area may be more informative in this model because mechanical CPR produces a rapidly changing waveform. Similar mean values can arise from waveforms with different peaks, widths, and recoil phases. The broader prone intrathoracic and arterial contours help explain why pressure-time area increased even when mean intrathoracic pressure changed little.

### Comparison with prior studies

Prior reports and reviews of prone CPR have described arterial pressures comparable with or higher than those generated in the supine position [1–4,8,9]. The present study reproduced that direction of effect under standardized mechanical compression and added synchronized pleural and peritoneal pressure measurements. The pattern is also consistent with established descriptions of thoracic and abdominal pump contributions to pressure generation during CPR [10–14].

This study examined pressure transmission in human anatomy rather than circulation. Arterial pressure generated in a cadaver does not establish blood flow, coronary perfusion pressure, organ perfusion, or the likelihood of return of spontaneous circulation. These results should not be interpreted as evidence that prone CPR is clinically superior. They offer a possible physiologic explanation for how prone compressions can generate substantial arterial pressure and support further study when immediate supination is not feasible.

### Limitations

This study included two biological specimens. Repeated trials improved the description of within-specimen consistency but did not create additional independent cadavers. The fixed-effects models used trial-level variation as the error term and were treated as exploratory. The small number of specimens also limited assessment of variation related to donor anatomy, body habitus, and tissue condition. Counterbalancing reduced systematic sequence bias, but one cadaver in each sequence was insufficient to evaluate carryover.

Fresh-frozen cadavers lack vascular tone, myocardial contraction, active venous return, gas exchange, autonomic regulation, and tissue perfusion [18,19]. Freeze-thaw changes may alter chest wall stiffness, abdominal compliance, vascular patency, and cavity behavior. Vascular conditioning reduced clot burden but could not restore living circulation or an intact capillary bed. The saline-based conditioning solution also may differ from blood in viscosity and hematologic behavior, which may have altered arterial waveform damping and transient pressure transmission. Additionally, the catheters measured local pleural and peritoneal pressure rather than pressure throughout each cavity.

The airway was clamped to standardize the thoracic pressure environment. Findings may differ with ventilation, airway leak, positive end-expiratory pressure, or an open airway [20,21]. Only one compression device and one prone compression region were studied. Diaphragm motion, chest displacement, blood flow, right atrial pressure, and coronary perfusion pressure were not measured. The proposed reduction in caudal pressure dissipation remains a mechanistic interpretation rather than a directly observed event and future studies involving fluoroscopy would be beneficial.

## Conclusions

In two fresh-frozen human cadavers, prone mechanical CPR produced higher arterial pressure, higher intra-abdominal pressure, and higher peak intrathoracic pressure than supine CPR. It also reversed the mean peritoneal-to-pleural pressure gradient and increased intrathoracic and arterial pressure-time area. Together, these findings in our pilot study suggest that abdominal counterpressure may limit caudal pressure dissipation and sustain a larger intrathoracic pressure pulse. Direct measurement of diaphragm motion and blood flow is needed to test this mechanism.

## Ethics approval and consent for anatomical donation

The UMCIRB determined that this cadaveric project was not human subjects research. Consent for anatomical donation and permitted research or educational use was managed by the body donation service before specimen release. No names, dates, record numbers, or other direct donor identifiers were included in the study dataset or manuscript.

## Data availability

The data supporting this study, including the Excel workbook and native LabChart recordings, are publicly available in Mendeley Data [22].

## Acknowledgments

The authors thank the anatomical donors and their families, the Middle Tennessee School of Anesthesia for laboratory access, and the Medical Education and Research Institute for coordinating specimen access. Anderson Funeral Home, Sean Johnson, and Katlyn Johnson provided technical assistance with specimen preparation.

## Declaration of AI

The authors declare that no generative artificial intelligence or AI-assisted technologies were used in the research process or in the preparation, writing, editing, or revision of this work. All content was developed and reviewed solely by the authors, who take full responsibility for its accuracy, integrity, and final form.

